# Norepinephrine controls the gain of the inhibitory circuit in the cerebellar input layer

**DOI:** 10.1101/567172

**Authors:** Frederic Lanore, Jason S. Rothman, Diccon Coyle, R. Angus Silver

## Abstract

Golgi cells (GoCs) are the main inhibitory interneurons in the input layer of the cerebellar cortex and are electrically coupled together, forming syncytia. GoCs control the excitability of granule cells (GCs) through feedforward, feedback and spillover-mediated inhibition. The GoC circuit therefore plays a central role in determining how sensory and motor information is transformed as it flows through the cerebellar input layer. Recent work has shown that GCs are activated when animals perform active behaviours, but the underlying mechanisms remain poorly understood. Norepinephrine (NE), also known as noradrenaline, is a powerful modulator of network function during active behavioral states and the axons of NE-releasing neurons in the *locus coeruleus* innervate the cerebellar cortex. Here we show that NE hyperpolarizes the GoC membrane potential, decreases spontaneous firing and reduces the gain of the spike frequency versus input-current relationship. The GoC membrane hyperpolarization can be mimicked with an α_2_-noradrenergic agonist, inhibited with a specific α_2_ antagonist and is abolished when G protein-coupled inwardly-rectifying potassium (GIRK) channels are blocked. Moreover, NE reduces the effective electrical coupling between GoCs through a persistent sodium current (I_NaP_)-dependent mechanism. Our results suggest that NE controls the gain of the GoC inhibitory circuit by modulating membrane conductances that act to reduce membrane excitability and decrease electrical coupling. These mechanisms appear configured to reduce the level of GoC inhibition onto GCs during active behavioural states.

## Introduction

The cerebellum is thought to play a key role in motor control by calculating errors in movements and learning to correct for them (De Zeeuw et al., 2011; Heiney et al., 2014; Herzfeld et al., 2018). It is also thought to predict the sensory consequences of active movement, thereby disentangling self-generated signals from external events (Cullen, 2012; Kennedy et al., 2014; Proville et al., 2014). Mossy fibres (MFs) convey sensory and motor information to the input layer of the cerebellar cortex (Arenz et al., 2008; Chen et al., 2017; Proville et al., 2014; Rancz et al., 2007; van Kan et al., 1993), where the two paths of information are combined and projected into a higher dimensional representation through a highly divergent circuit structure. This, together with decorrelation and sparsening of activity is thought to separate activity patterns, making downstream associative learning by Purkinje cells more effective (Albus, 1971; Billings et al., 2014; Cayco-Gajic et al., 2017; Litwin-Kumar et al., 2017; Marr, 1969; Tyrrell and Willshaw, 1992). Golgi cells (GoCs) are the main inhibitory neurons in the cerebellar input layer. They are thought to play a central role in pattern separation by providing strong inhibition onto granule cells (GCs) through feedforward, feedback and spillover-mediated inhibition (Brickley et al., 1996; Cesana et al., 2013; Crowley et al., 2009; Kanichay and Silver, 2008; Rossi et al., 2003). Nevertheless, whole-cell patch-clamp recordings have revealed that GCs can fire vigorously during locomotion (Powell et al., 2015) and calcium imaging of the GC population shows that a large fraction of GCs are activated when animals are performing active behaviors (Giovannucci et al., 2017; Knogler et al., 2017; Sylvester et al., 2017; Wagner et al., 2017). However, it is not known whether elevated GC activity is predominantly due to increased MF excitatory drive or neuromodulation-mediated disinhibition of GCs, as observed in other brain regions (Froemke et al., 2007; Fu et al., 2014; Letzkus et al., 2015; 2011; Poorthuis et al., 2014; Toth et al., 1997).

The activity of neurons that release neuromodulators such as norepinephrine (NE, or noradrenaline) often correlates with the behavioural state (McGinley et al., 2015; Reimer et al., 2014; 2016). The *locus coeruleus* (LC) is one of the main sources of NE in the brain and the activity of neurons within this nucleus increases with arousal and active behaviors (Joshi et al., 2016; Reimer et al., 2016). The cerebellum is innervated by noradrenergic inputs (Berger et al., 1979; Kimoto et al., 1981; Loughlin et al., 1986; Olson and Fuxe, 1971; Robertson et al., 2013) that arise from the LC (Kimoto et al., 1978). NE has been shown to modulate Purkinje cell activity (Freedman et al., 1977; Hoffer et al., 1973) via pre- and postsynaptic regulation of glutamate (Dolphin, 1982; Moises et al., 1981) and GABA signaling (Freedman et al., 1976; Waterhouse et al., 1982). NE also modulates excitatory synaptic transmission onto Purkinje cells as well as down regulating the induction of synaptic plasticity (Carey and Regehr, 2009). Moreover, enhanced glial cell activity during locomotion is mediated by noradrenergic pathways (Bazargani and Attwell, 2017; Paukert et al., 2014). While noradrenergic inputs are present in all of the layers of the cerebellar cortex (Berger et al., 1979; Olson and Fuxe, 1971), noradrenergic modulation has only been reported in the molecular layer, where most plasticity is thought to occur (Gao et al., 2012). But, it is not known whether noradrenergic modulation alters the activity of the cerebellar input layer during active behavioural states.

To investigate whether NE regulates activity in the cerebellar input layer, we performed whole-cell patch-clamp recordings from GoCs and GCs in acute slices of mouse cerebellar cortex. Application of low concentrations of NE hyperpolarized GoCs and reduced their spontaneous firing rate. Pharmacological approaches revealed that these effects were mediated by α_2_-adrenergic receptor activation of G protein-coupled inwardly-rectifying potassium (GIRK) channels, which are abundant on GoCs (Watanabe and Nakanishi, 2003). Moreover, application of NE reduced electrical coupling between GoCs. These results suggest that NE decreases the excitability of the inhibitory circuit in the input layer during active behavioural states.

## Results

### Origin of tyrosine hydroxylase-positive axons in mouse cerebellar cortex

Noradrenergic neurons express tyrosine hydroxylase (TH), an enzyme involved in the synthesis of NE (Nagatsu et al., 1964) that identifies catecholaminergic structures (Köhler and Goldstein, 1984). Previous studies have found diffuse TH-expressing fibres in rodent cerebellum (Berger et al., 1979; Bloom et al., 1971; Dietrichs, 1988; Felten et al., 1986; Landis et al., 1975; Olson and Fuxe, 1971; Pasquier et al., 1980). LC is known to be the main source of cerebellar afferents in rat (Kimoto et al., 1978; Olson and Fuxe, 1971). To investigate whether this is also the case in mice, we performed retrograde tracing with red fluorescent retrobeads injected in lobule IV/V of the cerebellar cortex and performed immunohistochemistry against TH in 3 mice. TH-expressing axons were distributed through the layers of cerebellum (Figure 1A) and retrobeads and TH-positive cells colocalized in LC (Figure 1B, C). Moreover, we found no trace of retrobeads in the Ventral Tegmental Area (VTA; Figure 1D, E) consistent with a previous study (Wagner et al., 2017). These experiments show that noradrenergic axons originating in LC project to lobule IV/V of mouse cerebellar cortex.

**Figure 1:**
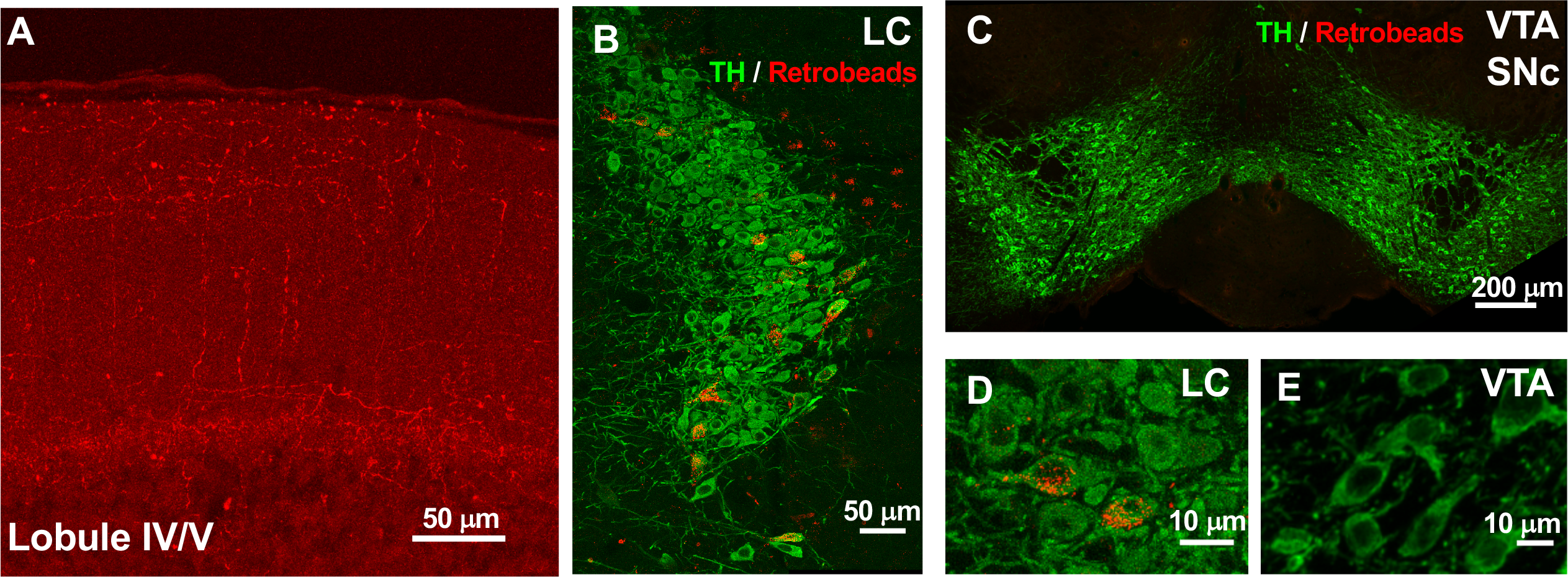
Noradrenergic axonal projections to Lobule IV/V of the the cerebellar cortex arise from the *locus coeruleus*. **A**, representative coronal section of lobule IV/V in the cerebellum showing a homogeneous distribution of immunolabeled TH-positive axons in red. **B**, sections through *locus coeruleus* (LC) showing TH immunoreactivity (green) and retrobeads (red) that had been injected in lobule IV/V of the cerebellar cortex and retrogradely transported to LC. C Higher magnification image showing accumulated of beads in the somatic compartments of LC. **D** and **E** as for B and C but for the Ventral Tegmental Area (VTA), where no retrobeads were found.

### Norepinephrine reduces Golgi cell firing rate and the gain of the frequency-current relationship

We next investigated the effect of NE on GoC activity in acute slices of cerebellar cortex from young adult mice (P22-33) by recording spontaneous action potential firing (Forti et al., 2006) with the non-invasive loose-cell-attached voltage-clamp technique (Figure 2A). GoCs in the cerebellar vermis spiked at an average rate of 7 Hz (range: 2 – 27 Hz, mean: 7 ± 1 Hz; n = 22), consistent with the firing rate recorded *in vivo* (Dugué et al., 2009; Hartmann and Bower, 1998; van Kan et al., 1993). Bath application of 10 μM NE for 5 min decreased the spontaneous firing rate of GoCs (Ctr: 7 ± 2 Hz, NE: 1 ± 2 Hz; Wilcoxon signed-rank test, p = 0.031, n = 6; Figure 2B), an effect that was largely reversed upon washout. To examine whether NE affects GoC excitability, we carried out whole-cell current-clamp recordings from GoCs during bath application of 10 μM NE. This induced a hyperpolarization of the resting membrane potential from - 56 ± 0.5 mV to −59 ± 0.8 mV (n = 14; Wilcoxon signed-rank test, p = 0.0002; Figure 2C, D) and larger depolarizing current pulses were required to reach spike threshold (Figure 2E, F). NE also reduced the slope (gain) of the relationship between firing frequency and injected current (*f*-*I* curve; Figure 2F-H). Quantification of these effects revealed that NE approximately halved the slope of the *f*-*I* curve (Ctr: 0.15 ± 0.01, NE: 0.08 ± 0.02; n = 8; Wilcoxon signed-rank test, p = 0.016; Figure 2G) and increased the rheobase (Ctr: 65 ± 8 pA, NE: 102 ± 17 pA; n = 9; Wilcoxon signed-rank test, p = 0.0078; Figure 2H). These results show that NE strongly modulates GoC excitability, altering both the gain of the *f*-*I* curve and the amount of current required to reach spike threshold.

**Figure 2:**
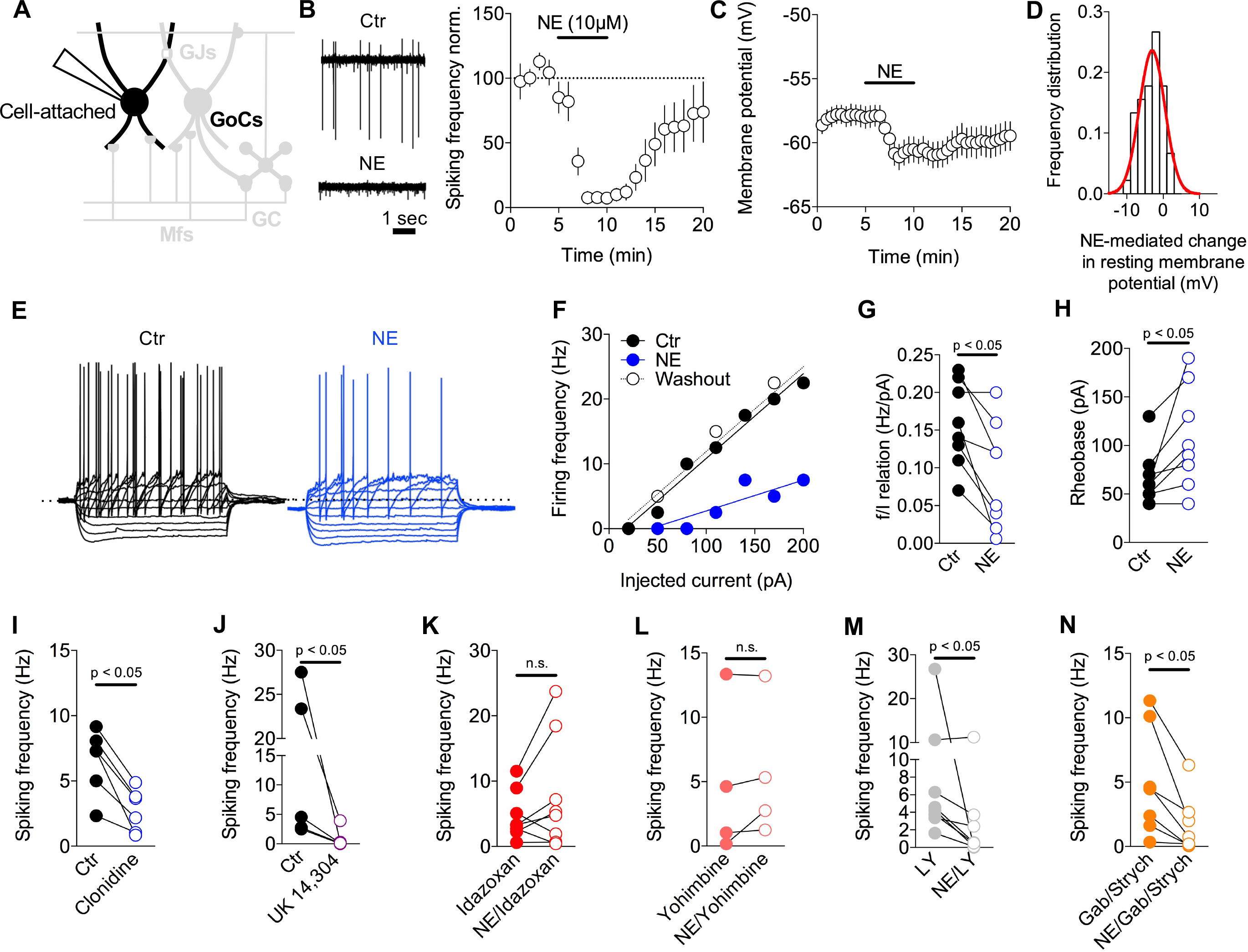
Norepinephrine decreases Golgi cell spontaneous firing via α_2_-adrenergic receptor subunit activation. **A**, Schematic of the GC layer showing a patch-clamp recording from a GoC. **B**, Example of spontaneous GoC firing recorded in cell-attached mode in control ACSF (top) and 10 μM NE (bottom). Summary graph (right) of the time course of spontaneous GoC firing before, during and after bath application of 10 μM NE (n = 6). Values are normalized to the first 5 min of control. **C**, Time course of GoC resting membrane potential, recorded in whole cell mode, before, during and after bath application of 10 μM NE (n = 14). **D**, Frequency distribution of the NE-mediated change in resting membrane potential (n = 45). **E**, Example GoC membrane voltage in control ACSF (left) and 10 μM NE (right) in response to 400-ms current steps from −200 to +200 pA. **F**, Relationship between spike frequency and injected current (f/I relationship) for the GoC in (**E**), including washout after NE. **G-H**, Summary of the effect of NE on the slope (**G**; n = 8) and rheobase (**H**; n = 9) of the GoC f/I relationship (paired t-test, * p < 0.05, ** p < 0.01). **I-J**, Effects of bath application of the α_2_-adrenergic receptor agonist Clonidine (**I**; 1 μM) and UK 14,304 (**J**; 15 μM) on GoC spontaneous firing rate. **K-L**, effect of NE on GoC firing rate in the presence of the α_2_-adrenergic receptor antagonists idazoxan (**K**; 1 μM) and Yohimbine (**L**; 15 μM). **M-N**, effect of NE on GoC firing rate in the presence of the mGluR antagonist LY341495 (**M**; 100 μM) or a combination of the GABAergic antagonist Gabazine and the glycinergic antagonist Strychnine (**N**; 10 and 0.5 μM, respectively).

### Modulation of Golgi cells is mediated by α_2_-adrenergic receptor activation

To identify the receptor subtype underlying the NE-mediated decrease in GoC excitability, we examined the effects of selective adrenergic agonists and antagonists. We focused on α_2_-adrenergic receptors as they are expressed in GoCs (Wang et al, 1996; Schambra et al, 2005). Application of the α_2_-adrenergic receptor agonist clonidine (1 μM) markedly reduced GoC spiking activity (Ctr: 6 ± 1 Hz, clonidine: 2 ± 0.6 Hz, n = 6; Wilcoxon signed-rank test, p = 0.031; Figure 2I). A similar suppression of firing rate was obtained with the α_2_-adrenergic receptor agonist UK 14,304 (15 μM; Ctr: 10 ± 4 Hz, UK 14,304: 0.7 ± 0.6 Hz, n = 6; Wilcoxon signed-rank test, p = 0.031; Figure 2J). Moreover, the α_2_-adrenergic receptor antagonist idazoxan (1 μM) blocked the NE-mediated inhibition of GoC firing (idazoxan: 4 ± 1 Hz, NE/idazoxan: 7 ± 3 Hz, n = 8, Wilcoxon signed-rank test, p = 0.1953, Figure 2K). A similar effect was produced by the α_2_-adrenergic receptor antagonist yohimbine (15 μM) (yohimbine: 4 ± 3 Hz, NE/yohimbine: 5 ± 2 Hz, n = 4, Wilcoxon signed-rank test, p = 0.25, Figure 2L). We also tested for α_1_-adrenergic receptors by applying the antagonist Prazonin (0.1 μm), but this did not block NE-mediated inhibition of GoC firing (Prazonin: 5 ± 5 Hz, NE/Prazonin: 0.1 ± 0.1 Hz, n = 5, Wilcoxon signed-rank test, p = 0.05; data not shown). This pharmacological profile suggests NE inhibits GoC spiking through an α_2_-adrenergic receptor-dependent mechanism.

While it seems likely that the α_2_-adrenergic receptors are located on the GoC membrane (Schambra et al., 2005; R. Wang et al., 1996) and could provide a direct mechanism of action for NE, an indirect mechanism could also exist. For example, NE activation of glial cells (Paukert et al., 2014), or another neuronal type, could result in glutamate, GABA or glycine release onto GoCs that could potentially mediate the membrane hyperpolarization observed in the presence of NE. Since mGluR2 receptors are expressed at high levels on GoC dendrites and can also hyperpolarize GoCs (Watanabe and Nakanishi, 2003), we tested whether they were involved in NE-mediated inhibition of spontaneous GoC firing. Bath application of LY341495 (100 μM), a potent group II mGluR antagonist, did not abolish the NE-mediated inhibition of GoC firing (LY: 8 ± 3 Hz, LY/NE: 2 ± 1 Hz, n = 6; Wilcoxon signed-rank test, p = 0.031; Figure 2M). To test whether GABA release from glial cells (Lee et al., 2010) or Lugaro cells (Dieudonné, 1995) was involved in NE-mediated inhibition of spontaneous GoC firing, we bath applied gabazine (10 μM) and strychnine (0.5 μM) to block GABAA and glycine receptors. However, these antagonists did not block the NE-mediated decrease in GoC firing rate (Gab/Strych: 4 ± 1 Hz, Gab/Strych/NE: 1 ± 0.8 Hz, n = 7; Wilcoxon signed-rank test, p = 0.016; Figure 2N). These results are all consistent with NE acting directly on α_2_-adrenergic receptors located on the GoC membrane.

### Norepinephrine-mediated hyperpolarization is mediated by GIRK channels

To investigate the origin of the ionic conductance(s) responsible for NE-mediated hyperpolarization and inhibition of firing in GoCs, we performed whole-cell voltage-clamp recordings at a holding potential of −60 mV (Figure 3A). Bath application of NE (10 μM) induced an outward current in GoCs (28 ± 6 pA, n = 24, Figure 3B, C) that was blocked by the α_2_-adrenergic antagonist idazoxan (NE: 29 ± 6 pA, measured over an 2 min window during NE application, immediately prior to antagonist application; NE+idazoxan: 5 ± 3 pA, measured over an 2 min window at the end of NE+antagonist application; n = 20; Wilcoxon signed-rank test, p < 0.0001; Figure 3D). This was not due to suppression of a tonically-activated outward current since bath application of idazoxan alone had no effect on the holding current (Figure 3D). Previous studies on neurons in the hypothalamus have shown that the α_2_-adrenergic subunit activates G protein-coupled inwardly-rectifying potassium (GIRK) channels (Li and van den Pol, 2005) and GIRK channels are highly expressed in GoCs (Watanabe and Nakanishi, 2003). To investigate whether this ionic mechanism underlies the NE-mediated outward current in GoCs, we applied the toxin Tertiapin-Q which selectively blocks GIRK channels (Jin and Lu, 1999; Malik and Johnston, 2017). Bath application of Tertiapin-Q (0.5 μM) blocked the NE-mediated outward current, confirming the involvement of GIRK channels (NE: 29 ± 6 pA, NE+Tertiapin-Q: 13 ± 4 pA; n = 7; Wilcoxon signed-rank test, p = 0.016; Figure 3E). We also examined the effect of BaCl_2_, which is known to block GIRK currents in GoCs (Watanabe and Nakanishi, 2003). Consistent with the effect of Tertiapin-Q, 100 μM BaCl_2_ abolished the outward current generated by 10 μM NE (NE: 22 ± 6 pA, NE+BaCl_2_: −2 ± 5 pA; n = 6; Wilcoxon signed-rank test, p = 0.031; Figure 3F). To test the involvement of G proteins in producing the NE-mediated outward current in GoCs, we replaced GTP in the patch pipettes with the non-hydrolyzable GTP analog GTPγS (1 mM), which confers constitutive G protein activity (Gonzalez et al., 2018; Li and van den Pol, 2005). Under these conditions application of NE did not induce an additional current (NE: −2 ± 3 pA from baseline; n = 12; Wilcoxon signed-rank test, p = 0.458; Figure 3G) confirming the involvement of G protein coupling in the NE-activated outward current. Taken together these results suggest that NE acts via α_2_-adrenergic receptors to activate GIRK channels in GoCs, which generate an outward current that hyperpolarizes the GoC membrane potential.

**Figure 3:**
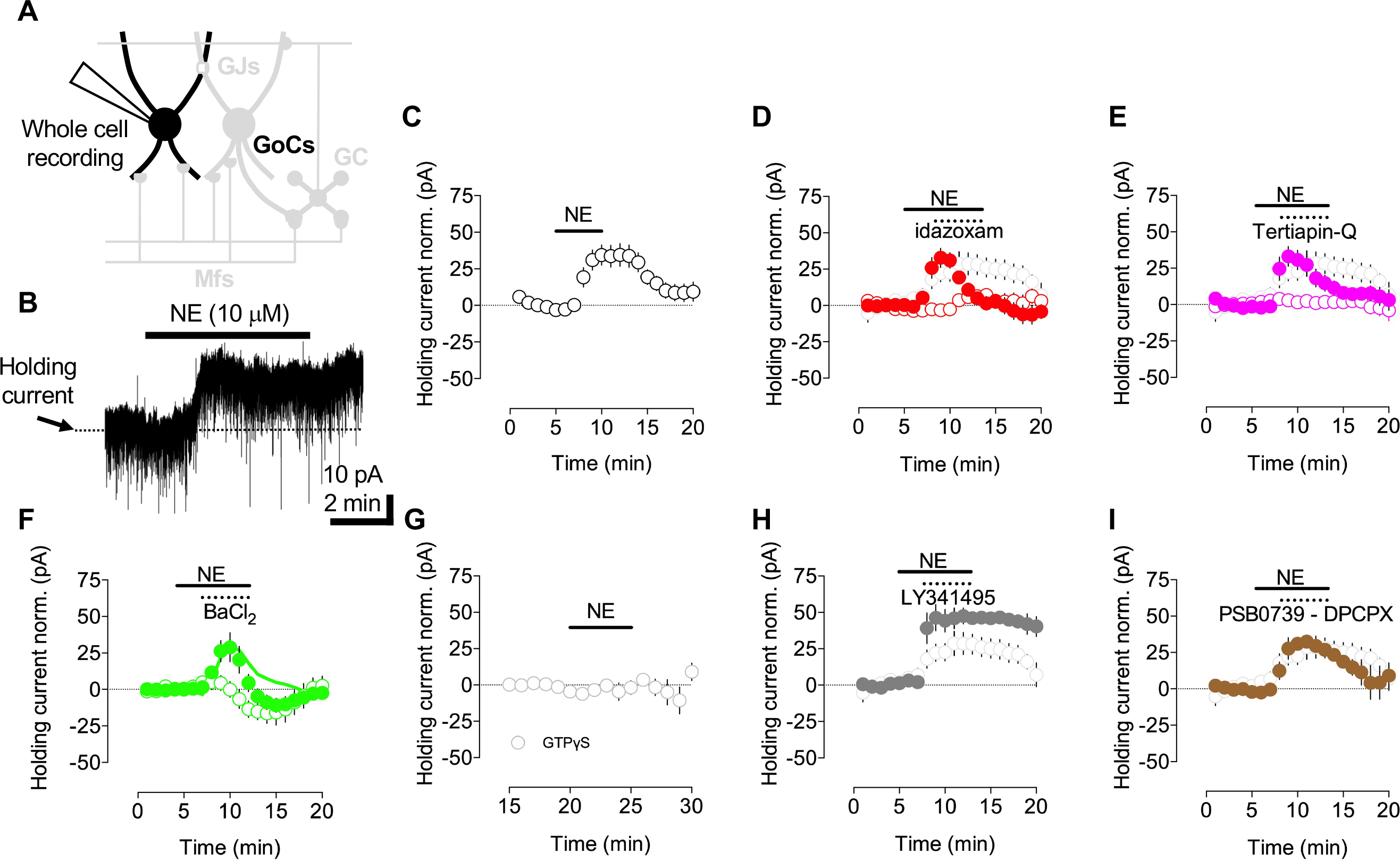
Norepinephrine activates an outward current in Golgi cells mediated by GIRK channels. **A**, Schematic of the GC layer showing a whole-cell voltage-clamp recording from a GoC. **B**, Bath application of 10 μM NE induced an outward current in a GoC. **C**, Summary of the NE-mediated outward current in 24 GoCs. **D**, Effect of the α_2_-adrenergic receptor antagonist idazoxan (1 μM) on the NE-mediated outward current (closed red circles). Bath application of idazoxan by itself (open red circles). Light gray circles in panel **D** to **I** denote bath application of 10 μM NE by itself (n = 4). **E**, Effect of Tertiapin-Q (0.5 μM, a specific inhibitor of GIRK channels) on the NE-mediated outward current (closed pink circles). Bath application of Tertiapin Q in the absence of NE (open pink circles) on the holding current. **F**, Effect of BaCl_2_ (100 μM) on the NE-mediated outward current in GoCs (closed green circles. Open green circles denote the GoC response to BaCl_2_ in the absence of NE. Green line denotes the difference between the green closed and open circles. **G**, Effect of NE on GoC current when GTP was replaced by GTPγS in the patch pipette. **H**, NE-mediated outward current in GoCs in the presence of LY341495 (100 μMa group II mGluR antagonist; grey filled circles). **I**, Effect of a cocktail of PSB 0739 (0.1 μM), a P2Y_12_ receptor antagonist, and DPCPX (100 nM), a specific antagonist for A1 adenosine receptors, on the NE-mediated outward current in GoCs.

Besides α_2_-adrenergic receptors, GIRK channels are known to be activated by mGluR2 receptors (Watanabe and Nakanishi, 2003), A1 adenosine receptors (Kim and Johnston, 2015; X. Wang et al., 2013) and purinergic receptors (Erb and Weisman, 2012; Filippov et al., 2004). We tested whether these signaling pathways contributed directly or indirectly (Bazargani and Attwell, 2017) to the NE-mediated GIRK activation in GoCs by bath applying the group II mGluR antagonist LY341495 (100μM; NE: 42 ± 9 pA, NE+LY: 46 ± 3 pA; n = 6; Wilcoxon signed-rank test, p = 0.688; Figure 3H) and a cocktail of the P2Y12 purinergic receptor antagonist PSB 0739 (0.1 mM; (Madry et al., 2018)) and A1 adenosine receptor antagonist DPCPX (100 nM; NE: 23 ± 5 pA, NE+PSB/DPCPX: 26 ± 3 pA; n = 7; Wilcoxon signed-rank test, p = 0.938; Figure 3I). None of these antagonists affected the NE-mediated outward current, consistent with NE activating GIRK channels via α_2_-adrenergic receptors on the GoC membrane.

### Effect of norepinephrine on excitatory synaptic transmission onto Golgi cells

NE has previously been shown to reduce the probability of glutamate release at climbing fibre to Purkinje cell connections (Carey and Regehr, 2009). To investigate whether NE also reduces the excitatory drive onto GoCs, we examined its effect on EPSCs arising from MFs and parallel fibres (PFs). Putative MF EPSCs were recorded from GoCs by stimulating in the cerebellar white matter tract (Kanichay and Silver, 2008) (Figure 4A). Bath application of 10 μM NE had no effect on the EPSC amplitude (Ctr: 57 ± 6 pA, NE: 51 ± 6 pA, Washout: 57 ± 9 pA, n = 10, Wilcoxon signed-rank test, p = 0.232), paired-pulse ratio (Ctr: 0.99 ± 0.07, NE: 0.94 ± 0.07, Washout: 1 ± 0.09, n = 10, Wilcoxon signed-rank test, p = 0.275) or the short term plasticity induced by a 5-pulse train at 100 Hz (amplitude ratio EPSC5/EPSC1: Ctr: 0.9 ± 0.06, NE: 0.87 ± 0.09, Washout: 1 ± 0.13, n = 10, Wilcoxon signed-rank test, p = 0.49) (Figure 4B).

**Figure 4:**
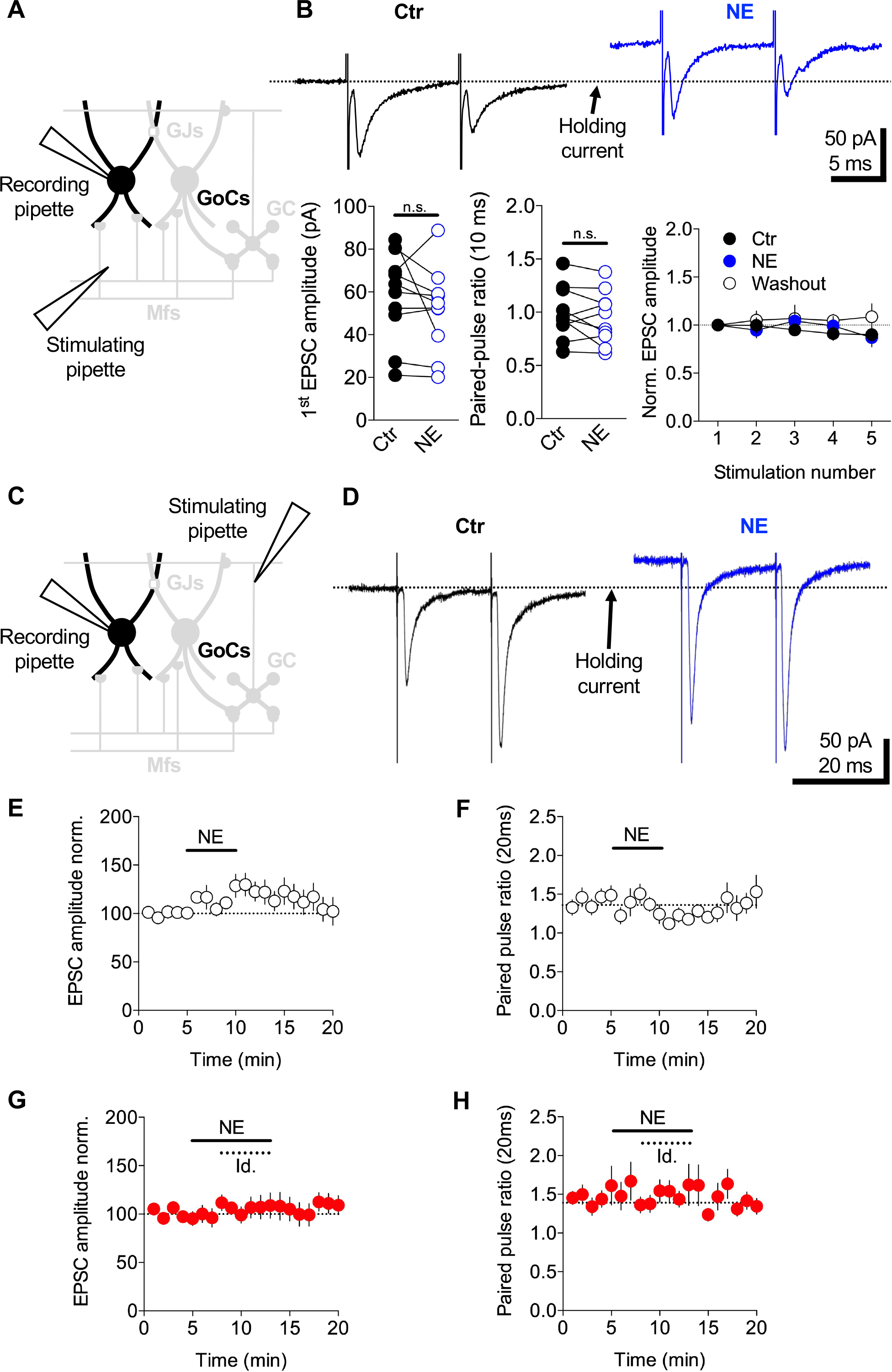
Norepinephrine has little effect on excitatory synaptic transmission. **A**, Schematic of the cerebellar cortex showing whole-cell voltage-clamp of a GoC while MF inputs were stimulated via an electrode in the white matter. **B**, Mean MF EPSCs recorded from a GoC in response to 5 stimuli delivered at 100 Hz (only first two EPSCs are shown) in control ACSF (black) and 10 μM NE (blue). Bath application of 10 μM NE had no significant effect on the amplitude of MF-GoC synaptic transmission or the short-term plasticity. Note the holding current change during NE application. **C**, Same as (A) except the stimulating electrode was placed in the molecular layer to trigger synaptic release from PFs. **D**, Mean PF EPSCs recorded from a GoC in response to 2 stimuli delivered at 50 Hz in control ACSF (black) and 10 μM NE (blue). **E-F**, Summary graphs showing the effect of NE on the normalized amplitude of the first EPSC (left) and paired-pulse ratio (right). **G-H**, In the presence of the α_2_-noradrenergic receptor specific antagonist idazoxan (1 μM) NE had no significant effect on the EPSC amplitude or the paired-pulse ratio.

We next examined the effect of NE on PF evoked EPSCs, recorded from GoCs by stimulating PFs in the molecular layer (Figure 4C). In contrast to MF-EPSCs, application of NE produced a modest but reliable increase in PF-GoC synaptic transmission (Ctr: 117 ± 21 pA, NE: 142 ± 26 pA, Washout: 124 ± 20 pA, n = 14; Wilcoxon signed-rank test, p = 0.009) (Figure 4D, E). This is likely due to an increase in presynaptic glutamate release as the paired-pulse ratio decreased during NE bath application compared to control (Ctr: 1.3 ± 0.06, NE: 1.2 ± 0.06, Washout: 1.2 ± 0.09, n = 14; Wilcoxon signed-rank test, p = 0.020) (Figure 4F). Both NE-mediated potentiation of the PF-GoC EPSC (Ctr: 149 ± 17, NE+idazoxan: 162 ± 25, Washout: 167 ± 28, n = 14; Wilcoxon signed-rank test, p = 0.5416; Figure 4G) and the decrease in paired-pulse ratio were abolished by the α_2_-adrenergic antagonist idazoxan (Ctr: 1.3 ± 0.05, NE+ idazoxan: 1.4 ± 0.06, Washout: 1.3 ± 0.06, n = 14; Wilcoxon signed-rank test, p = 0.952; Figure 4H). Thus, NE has no detectable effect on feedforward excitatory synaptic inputs from MFs, but it enhances feedback excitation from GCs.

### Modulation of Golgi cell electrical coupling by norepinephrine

GoCs are electrically coupled to their neighbors via gap junctions on their dendrites (Dugué et al., 2009; Szoboszlay et al., 2016; Vervaeke et al., 2010; 2012). Electrical coupling increases the gain of GoC networks by enabling excitatory synaptic charge to be shared across neighboring cells (Vervaeke et al., 2012). To investigate whether NE modulates the gain of the GoC network by altering electrical coupling, we performed whole-cell current-clamp recordings from pairs of GoCs (Szoboszlay et al., 2016; Vervaeke et al., 2010) (Figure 5A). Electrical coupling strength was quantified using the coupling coefficient (CC), which is the ratio between pre- and postsynaptic voltage changes during long current injections (400 ms and 100 pA) into the presynaptic GoC (Figure 5B). In control conditions with artificial cerebrospinal fluid (ACSF), the CC was 0.12 ± 0.01 (range: 0.04 – 0.23; n = 7 pairs), similar to previous findings (Vervaeke et al., 2010; Szoboszlay et al., 2016). We measured the CC every 20 seconds over a 5 min control baseline, a 5 min application of NE, and a washout (Figure 5B, C). Application of 10 μM NE markedly decreased the CC (Ctr: 0.14 ± 0.01, NE: 0.10 ± 0.01, washout: 0.13 ± 0.01; Wilcoxon signed-rank test, p = 0.0001; n = 7 pairs; Figure 5C, D). This effect was associated with a hyperpolarization of the resting membrane potential of the recorded GoC pairs (Ctr: −56 ± 0.6 mV, NE: −59 ± 0.9 mV; Wilcoxon signed-rank test, p = 0.009; n = 14; Figure 5B) and a decrease in the input resistance (R_input_; Ctr: 112 ± 8 MΩ, NE: 103 ± 8 MΩ; Wilcoxon signed-rank test, p = 0.004; n = 14; data not shown). To test whether the decrease in the CC arose solely from the decrease in R_input_, we calculated the apparent coupling conductance (Fortier and Bagna, 2006). This revealed that 10 μM NE induced a significant decrease in the apparent coupling conductance (Ctr: 1.4 ± 0.2 nS, NE: 1.2 ± 0.2 nS; Wilcoxon signed-rank test, p = 0.0009; n = 14; Figure 5C, E). These results indicate the reduction in the CC induced by NE arises from a decrease in both R_input_ and the apparent coupling conductance.

**Figure 5:**
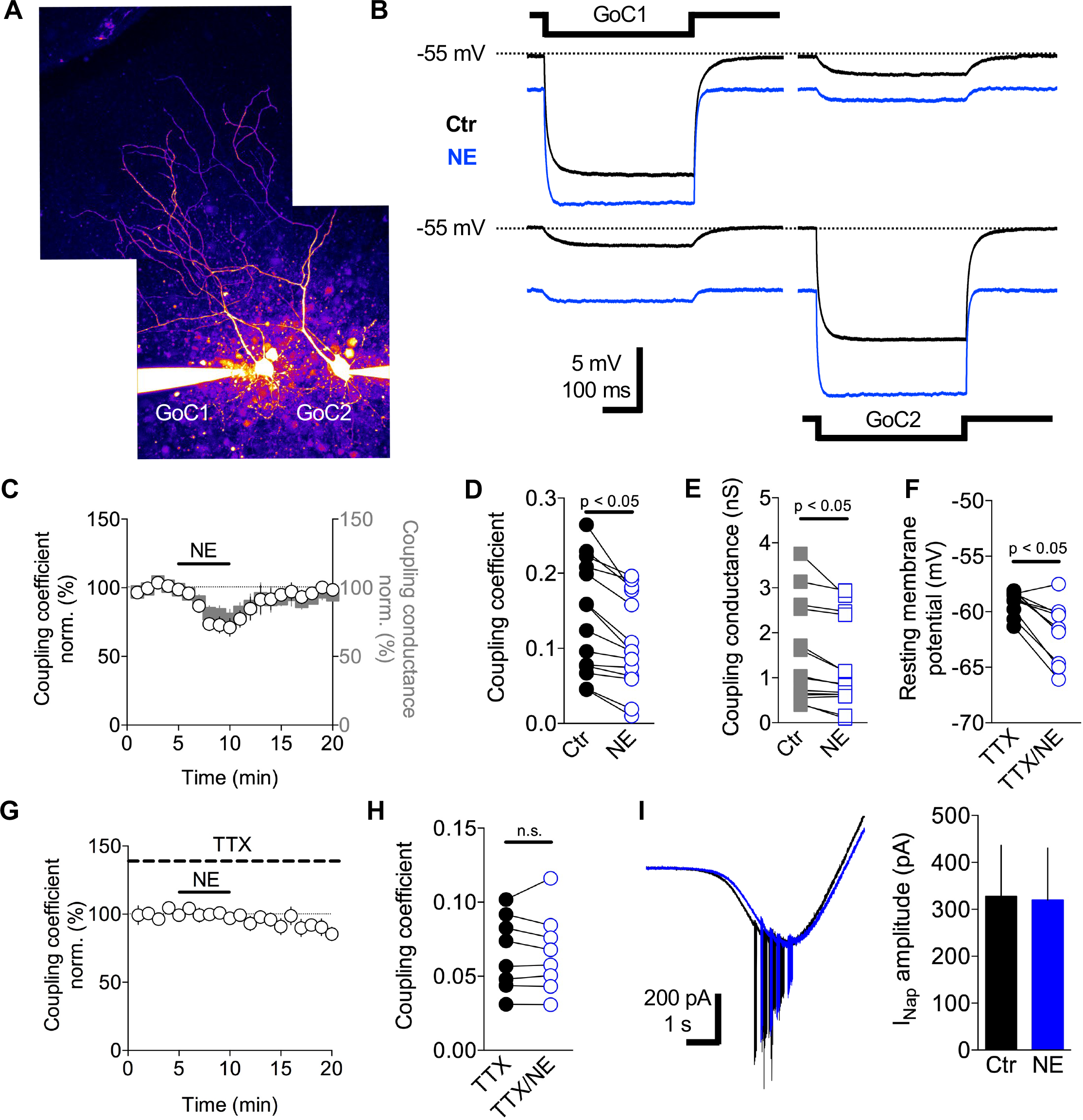
Norepinephrine-reduces electrical coupling between Golgi cells. **A**, Two-photon maximal projection image of two electrically coupled GoCs loaded with Alexa 594 (50 μM) via the patch pipettes. **B**, Membrane voltage recordings from the GoC pair in (**A**) in response to an injected current pulse (−100 pA) in GoC1 (top) and GoC2 (bottom) in control ACSF (black) and 10 μM NE (blue). **C**, After a 5 min baseline NE was bath applied (10 μM black bar) and the coupling coefficient (CC, white circles) and effective coupling conductance (gray squares) were monitored. **D-E**, Summary graphs showing the effect of NE (10 μM) on the CC and effective coupling conductance. **F**, Effect of NE (10 μM) on the membrane potential in the presence of TTX to block I_NaP_, NE (10 μM). (**G-H**) Effect of NE on the CC in the presence of TTX. **I**, Electrophysiological recordings of isolated I_Nap_ current in control (black trace) and in the presence of NE (10 μM, blue trace). Summary graph of the effect of NE on the amplitude of sodium current.

The apparent coupling conductance measured at the soma combines the conductance of the gap junctions (GJs), axial dendritic resistance (Szoboszlay et al., 2016) and any active processes present between the dendritic GJs and the soma. The persistent sodium current (I_NaP_) present in the perisomatic region of GoCs (Vervaeke et al., 2012) introduces a voltage dependence to GJ potentials in GoCs (Dugué et al., 2009). Depolarization promotes the opening of more I_NaP_ channels leading to the amplification of electrical coupling, while hyperpolarization induces the inverse effect (Dugué et al., 2009; Haas and Landisman, 2011). To investigate whether the NE-mediated modulation of the apparent coupling conductance was dependent on I_NaP_, we measured the CC and apparent coupling conductance in slices incubated with TTX (1 μM). In this condition, bath application of NE (10 μM) induced a modest hyperpolarization of the resting membrane potential of GoCs (TTX: −59 ± 0.3 mV, TTX/NE: −61 ± 0.8 mV; Wilcoxon signed-rank test, p = 0.004; n = 10; Figure 5F) similar to that in control ACSF. However, the CC between GoC pairs was unchanged during application of NE 10 μM (TTX: 0.06 ± 0.018, TTX/NE: 0.06 ± 0.01; Wilcoxon signed-rank test, p = 0.722; n = 5 pairs; Figure 5G, H) as was the apparent coupling conductance (TTX: 0.6 ± 0.1 nS, TTX/NE: 0.6 ± 0.1 nS; Wilcoxon signed-rank test, p = 0.547; n = 5 pairs; data not shown). Quantification of the sodium conductance present in GoCs, using a voltage clamp ramp protocol under conditions where GABAA, glycinergic, AMPA and NMDA receptors were pharmacologically blocked (**Methods**), did not reveal any significant change in the current amplitude (Ctr: 328 ± 108 pA, NE: 319 ± 111 pA, n = 6; Wilcoxon signed-rank test, p = 0.44; Figure 5I). These results suggest that the NE-mediated decrease in the apparent coupling conductance arises from an interplay between GIRK-mediated hyperpolarization and the persistent sodium conductance. NE-induced activation of the GIRK conductance hyperpolarizies the membrane, which turns off I_NaP_, reducing the effective coupling conductance and the gain of the GoC syncytium.

### Effect of norepinephrine on inhibitory synaptic transmission in the granule cell layer

GoC-mediated phasic release of GABA, spillover and tonic inhibition (Brickley et al., 1996; Crowley et al., 2009; Rossi et al., 2003) provide the main source of inhibition onto GCs. To examine whether NE affects inhibitory synaptic transmission onto GCs, in addition to modulating GoC gain, we performed GoC-GC paired recordings in the presence of D-AP5 (50 μM) and NBQX (10 μM) to block NMDA and AMPA receptors. Bursts of 20 action potentials at 20 Hz were triggered in GoCs by injecting brief depolarizing current steps, and IPSCs were simultaneously recorded in GCs (Figure 6A). Bath application of NE had no effect on R_input_ of GCs (Ctr: 1061 ± 94 MΩ, NE: 1031 ± 114 MΩ, n = 10; Wilcoxon signed-rank test, p = 0.56) or on GoC-GC inhibitory synaptic transmission, as assayed from the charge per second (Ctr: −4.4 ± 0.5 pA.s^−1^, NE: −3.2 ± 0.7 pA.s^−1^, washout: −3.9 ± 0.5 pA.s^−1^, n = 11; Wilcoxon signed-rank test, p = 0.32; Figure 6B). While the effect of NE on individual GoC-GC pairs was highly variable, the relationship between the change in GoC membrane potential following application of NE and the change in GC charge modulation exhibited a significant positive correlation (r = 0.760, p = 0.007; Figure 6C). NE-mediated hyperpolarization of GoCs was accompanied by a reduction in the total charge of IPSCs recorded from GCs. Phasic IPSCs and the slow component of the evoked inhibitory current, which we define as spillover-mediated (**Methods**; (Rossi et al., 2003)), were equally affected by NE (Phasic: r = 0.6924, p = 0.018; spillover: r = 0.7558, p = 0.007) (Figure 6D). Moreover, a negative modulation of the spillover inhibition was accompanied by a negative modulation of the fast IPSCs and vice versa (linear regression: R^2^ = 0.72, p = 0.001) (Figure 6E). To investigate whether these NE-mediated effects arose from analog signalling along the axon we mimicked the NE-induced hyperpolarization of the GoC membrane potential by injecting current (−9 ± 1 mV, range −15 to −6 mV; Ctr: −2.7 ± 0.7 pA.s^−1^, hyp: −2.9 ± 0.8 pA.s^−1^, n = 5; Wilcoxon signed-rank test, p = 0.63), by lowering external [K^+^] to 0.5 mM (−2 ± 1 mV; Ctr: −5.7 ± 1.7 pA.s^−1^, low [K^+^] −6.8 ± 2.3 pA.s^−1^, n = 4; Wilcoxon signed-rank test, p = 0.38) and by directly activating GIRK channels with the selective agonist ML297 10 μM (−6 ± 0.9 mV; Ctr: −3.9 ± 0.7 pA.s^−1^, ML297: −5 ± 1.1 pA.s^−1^, n = 4; Wilcoxon signed-rank test, p = 0.13). While it is possible that changes in GABA release following these perturbations was too subtle to detect, the fact that none of these perturbations significantly modulated the inhibitory charge in GCs, suggests that other mechanisms are responsible for the NE-dependent modulation of inhibition.

**Figure 6:**
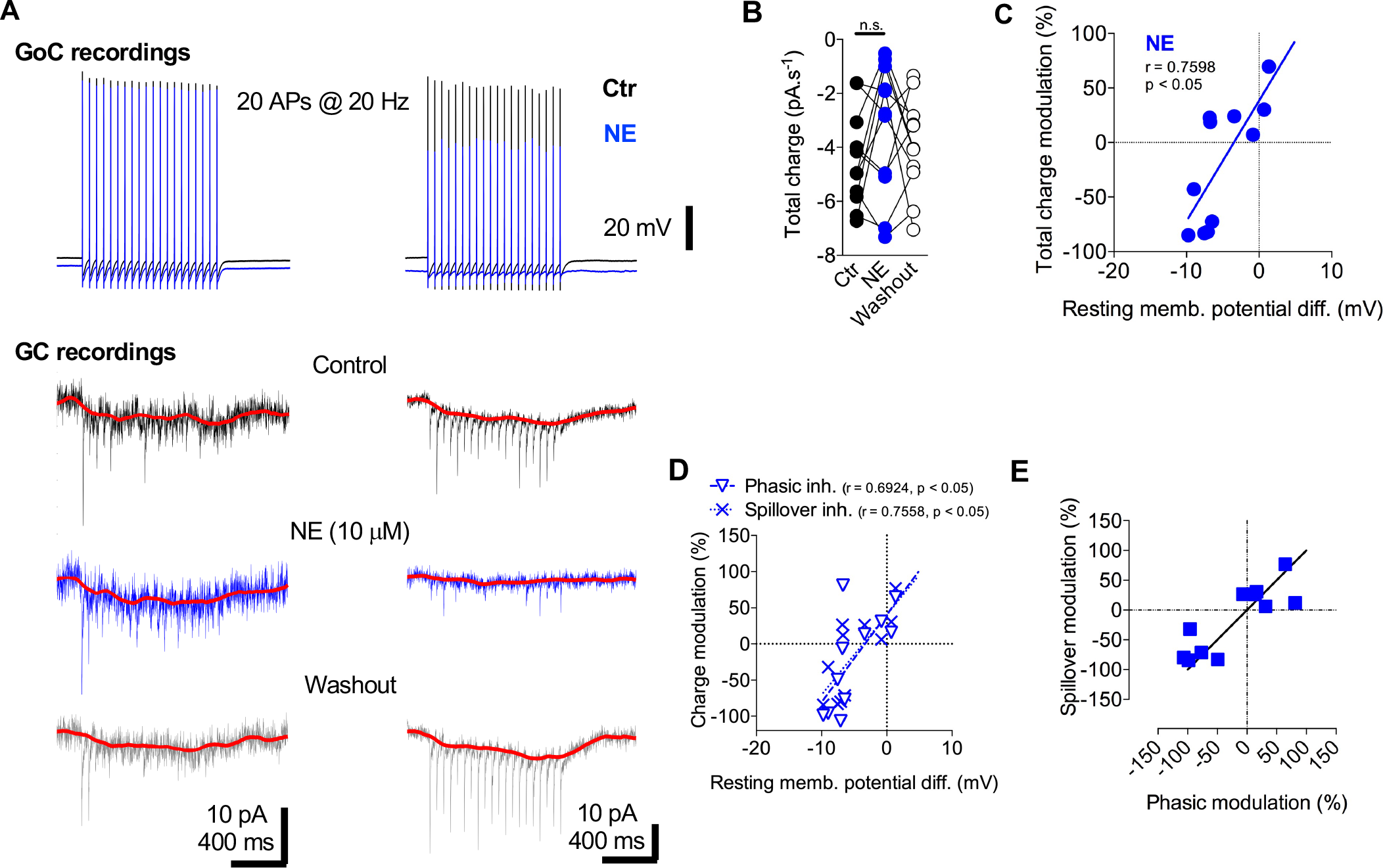
Norepinephrine modulates evoked postsynaptic inhibitory current onto granule cells. **A**, 20 action potentials at 20 Hz triggered with brief positive current injections via the somatic recording pipette in GoCs (top traces) and IPSCs recorded from GCs at a holding potential of −70 mV with a high chloride internal solution (bottom traces) in the presence of NBQX (10 μM) and D-AP5 (25 μM). NE (10 μM) was bath applied for 5 min (blue traces) after a 5 min baseline period (black traces). A low-pass filtered version (used for illustration purposes) of the slow spillover IPSC trace is shown overlaid with the total trains (red lines). The red line reflects the slow GABA spillover-mediated component of the IPSCs after subtraction of the phasic inhibition from the total IPSC. The figure shows two different paired recordings. On the example on the left, NE (10 mM) had no effect on the evoked inhibitory synaptic transmission while on the example on the right, NE inhibited the synaptically evoked responses. **B**, Summary graph of the total IPSC charge during baseline (Ctr), NE application and after a 10 min washout period. **C**, Plot of the total charge modulation (%) in function of the difference in resting membrane potential (mV) during NE bath application (blue dots). **D**, Size of modulation of synaptic charge mediated by phasic and spillover components as a function of the modulation in somatic membrane potential. **E**. relationship between the moulation in the phasic and tonic inhibitory components in granule cells.

## Discussion

In this study, we show that norepinephrine (NE) modulates the gain of GoCs, the main inhibitory interneurons in the input layer of the cerebellar cortex. NE hyperpolarizes membrane potential and reduces the spontaneous firing of GoCs by binding to α_2_-adrenergic receptors, which activate a powerful GIRK conductance. Moreover, NE also reduces electrical coupling between GoCs by lowering input resistance and by hyperpolarizing their membrane potentials, thereby reducing the boosting effects of the persistent sodium current (I_NaP_) on electrical signalling. By contrast, NE had little effect on excitatory and inhibitory chemical synaptic transmission. Our findings suggest that NE reduces the gain and increases the spike threshold of the Golgi cell network making it less sensitive to excitatory drive. These mechanisms could enable NE-releasing locus coeruleus (LC) projections to dynamically reduce inhibition in the GC layer during active behaviors.

### Mechanisms underlying norepinephrine-mediated modulation of Golgi cells

Several of our experimental observations indicate that NE modulates GoC excitability by activating α_2_-adrenergic receptors that turn on a GIRK conductance: i) GoC hyperpolarization is mimicked by α_2_-adrenergic receptor agonists and blocked by α_2_-adrenergic receptor-specific antagonists, ii) the NE-mediated outward current is blocked by the GIRK channel antagonists BaCl_2_ and Tertiapin-Q and iii) fully activating G-protein signaling occluded further GIRK channel activation by NE. The simplest explanation of these results is that NE directly binds to α_2_-adrenergic receptors on GoCs and activates a GIRK conductance. Indirect activation of glial cells by NE (Paukert et al., 2014) seems unlikely given that NE-mediated GoC hyperpolarization was unaffected when the common glial-neuron and neuron-neuron signalling via GABA, glutamate or adenosine (Bazargani and Attwell, 2017) was blocked. Moreover, reduction of GoC neuronal excitability by NE is consistent with the effects found on other types of neurons in the central nervous system (Arima et al., 1998; Carr et al., 2007; Kuo and Trussell, 2011; Li and van den Pol, 2005; North and Yoshimura, 1984; Williams and North, 1985; Xiao et al., 2009). Thus, our results indicate that NE modulates the gain of individual GoCs by binding α_2_-adrenergic receptors activating a powerful GIRK conductance.

Our paired recordings from GoCs show that the NE-induced GIRK conductance also modulates the electrical coupling between GoCs. Lowering the membrane resistance reduces electrical coupling because it reduces the voltage change induced by current flowing through gap junctions (Devor and Yarom, 2002). Moreover, GoCs express a persistent sodium conductance that boosts gap junction mediated signaling (Dugué et al., 2009; Haas and Landisman, 2011). NE-induce hyperpolarization reduces this boosting effect by shifting the operating voltage to a region where little of the persistent sodium conductance is active. These mechanisms are distinct from the direct modulation of gap junctional conductance at Goldfish Mauthner synapses (Pereda et al., 1998) and between interneurons in the neocortex (Landisman and Connors, 2005), which appear to operate on longer timescales. It is interesting to note that both of the effects of NE on GoCs lower the effective electrical coupling, which reduces the gain of electrically coupled networks to chemical excitatory input (Vervaeke et al., 2012).

### Effect of norepinephrine on synaptic transmission

Our results suggest that NE has rather subtle effects on excitatory synaptic drive onto GoCs and on GoC-mediated inhibition onto GCs. This is surprising given its widespread effects on synaptic transmission: NE downregulates synaptic transmission (Carey and Regehr, 2009; Delaney et al., 2007; Ohshima et al., 2017; H. X. Wang et al., 2013) and synaptic plasticity (Bear and Singer, 1986; Liu et al., 2017; Pettigrew and Kasamatsu, 1978) in different brain areas via presynaptic modulation of release probability (for review see: (Sara, 2009)). In the cerebellum, α_2_-adrenergic receptor activation at climbing fiber to Purkinje cell synapses inhibits excitatory synaptic transmission via a decrease in presynaptic release and impacts the induction of associative plasticity (Carey and Regehr, 2009). By contrast, application of NE had no detectable effect on excitatory synaptic transmission at MF to GoC synapses. However, α_2_-adrenergic receptor activation did lead to a modest increase in synaptic transmission and a decrease in the paired-pulse ratio at PF to GoC synapses, consistent with an increase in release probability. This is unusual as α_2_-adrenergic receptors are associated with downregulation of synaptic transmission, for example, via inhibition of voltage-gated Ca^2+^ channels (Bean, 1989; Dunlap and Fischbach, 1981). However, it is possible that this small effect is mediated by an indirect path involving glial cells in the molecular layer (Paukert et al., 2014).

Our GoC-GC paired recordings revealed that modulation of the inhibitory synaptic currents by NE was correlated with the magnitude of the hyperpolarization. Although this might suggest analog modulation of spike-evoked synaptic transmission (Debanne, 2004; Debanne et al., 2011) we could not mimic this effect by artificially hyperpolarizing the GoC membrane potential, by lowering external [K^+^] or by activating GIRK channels present on GoCs. These results suggest that NE-modulates release through an alternative mechanism, possibly a direct effect on GABA release through the same G protein pathway that activates the GIRK conductance.

### Origins and properties of norepinephrine-releasing fibres in the cerebellar cortex

Our tracing experiments suggest that TH expressing fibres found in lobules IV/V of mouse cerebellar cortex arise from the LC. This is consistent with previous work in rats, which also found that LC rather than VTA were the source of TH expressing fibres in the cerebellar cortex (Kimoto et al., 1978; Olson and Fuxe, 1971). Interestingly, LC also receives feedback projections from the cerebellum (Breton-Provencher and Sur, 2019) from Purkinje cell axons projections (Schwarz et al., 2015) forming a closed loop. The activity of neurons in LC has been shown to correlate strongly with active behaviors such as whisking and locomotion (Reimer et al., 2016; 2014). LC activation also correlates strongly with pupil diameter, which has also been associated with attention (McGinley et al., 2015; Reimer et al., 2016). These observations suggest that LC activity and thus NE release in the cerebellum will increase when animals are active. LC neurons are also thought to act as novelty detectors as NE release increases in response to novel sensory stimuli and with stimulus saliency (Grant et al., 1988; Takeuchi et al., 2016; Vankov et al., 1995).

### Implications of norepinephrine-mediated modulation of electrical coupling

GoCs are strongly electrically coupled through gap junctions on their dendrites (Dugué et al., 2009; Szoboszlay et al., 2016; Vervaeke et al., 2010). Electrical signalling is augmented by persistent sodium conductances, which amplify synaptic potentials as it flows to the soma (Dugué et al., 2009; Haas and Landisman, 2011). Electrical coupling tends to synchronize the firing of cells in a network (Connors and Long, 2004), but can also facilitate network desynchronization when synaptic excitation is sparse and gap junction potentials are dominated by inhibitory afterhyperpolarization, as found in GoCs (Vervaeke et al., 2010). Dendritic gap junctions can also redistribute synaptic charge, making networks of spontaneously active interneurons more sensitive to excitatory drive (Vervaeke et al., 2010). Our results showing that NE hyperpolarizes GoCs and reduces their effective electrical coupling is therefore likely to alter the properties of the GoC network in different ways.

First, the sensitivity or gain of the network to excitatory drive is expected to fall (Vervaeke et al., 2012), as synaptic charge shared with neighboring cells through dendritic gap junctions is less able to drive neighboring cells. This will make synaptic excitation of the inhibitory network less effective and could preferentially weaken feedback excitation via PF inputs, which are electrotonically more remote than MF inputs. These changes are expected to reduce the gain of the inhibitory network in the cerebellar input layer, which may enable the network to keep within the limited dynamic range of GoC firing during the elevations in excitatory drive that occur during active behaviours.

Second, NE-modulation of coupling between GoCs is expected to alter spike synchrony. When animals are in a quiet wakeful state, input to the cerebellar cortex and LC activity are both expected to be low. Under these conditions, intrinsic activity and strong electrical coupling is expected to promote spike synchronization in the GoC network, as observed under anesthesia (van Welie et al., 2016). Previous work has shown that electrically-coupled GoC networks contribute to local field potential (LFP) oscillations in the theta band (~ 8 Hz) (Dugué et al., 2009; Robinson et al., 2017). Indeed, such oscillations are present in the GC layer of awake and alert animals (Dugué et al., 2009; Hartmann and Bower, 1998; O’Connor et al., 2002). However, during active behaviours, elevated excitatory drive and reduced electrical coupling due to NE release from LC axons is expected to reduce neuronal synchronization. While speculative, this hypothesis is supported by the observation that GC-layer theta band oscillations disappear when rats display visible movements (Hartmann and Bower, 1998). Although activity-dependent cessation of theta oscillations could have several origins (e.g. (Vervaeke et al., 2010)), a decrease in GoC electrical coupling by NE is well placed to contribute to this phenomenon.

### Implications for sensorimotor processing and learning in the cerebellar cortex

GoCs provide strong inhibition to GCs through a tonic GABAA-mediated conductance, phasic release and spillover (Crowley et al., 2009). Tonic inhibition changes the gain of the GC spiking activity (Chadderton et al., 2004; Crowley et al., 2009; Hamann et al., 2002; Mitchell and Silver, 2003; Rothman et al., 2009) while phasic inhibition regulates GC spike timing precision (Duguid et al., 2015; Nieus et al., 2014). These inhibitory mechanisms are thought to play a key role in setting the high threshold for GC activation and sparsening population activity as signals flow from MFs to GCs (Billings et al., 2014). Indeed, strong inhibition and sparse coding has long been thought to play a key role in pattern separation prior to associative learning (Albus, 1971; Cayco-Gajic et al., 2017; Marr, 1969). But recent whole-cell recordings from GCs (Chen et al., 2017; Powell et al., 2015) and imaging of GC population activity in awake mice (Giovannucci et al., 2017; Knogler et al., 2017; Sylvester et al., 2017; Wagner et al., 2017) has reported high levels of activity during behaviour, challenging the idea that sensorimotor information is encoded as a sparse code (Cayco-Gajic and Silver, 2019). Moreover, recent theoretical work suggests that the input layer can perform pattern separation in non-sparse regimes, through mixing, expansion and decorrelation of inputs (Cayco-Gajic et al., 2017), which increase the dimensionality of the representation (Cayco-Gajic and Silver, 2019). Our results suggest that NE release from LC neurons could play a key role in elevating GC activity during active behavioural states, by lowering the gain of the inhibitory GoC network and dampening the input activity dependent increase in inhibitory tone. Enabling higher GC firing rates may help conserve information transmission through the input layer, since overly sparse codes lead to information loss (Billings et al., 2014).

Several studies have implicated the noradrenergic system in motor learning (Capasso et al., 1996; Keller and Smith, 1983; Mason and Iverson, 1974; McCormick and Thompson, 1982; Pompeiano, 1998; Watson and McElligott, 1984). High levels of GC activity during behaviour could facilitate plasticity at different locations in the cerebellar cortex (Gao et al., 2012). These include long-term potentiation of MF-GC synaptic transmission and the long-term increase in excitability of GC intrinsic membrane properties (Armano et al., 2000; Nieus et al., 2006). During active states when NE is likely to be released (Reimer et al., 2016), reduced GoC activity could aid the transfer of information from MFs to GCs, improving the reliability of firing at PF-Purkinje cell synapses, facilitating long-term plasticity and speeding learning. This hypothesis is supported by the fact that learning in the cerebellum is facilitated by locomotion (Albergaria et al., 2018) which is known to be correlated to an increase in arousal and the release of NE (Reimer et al., 2016). Our results showing that NE has a powerful effect on the gain of the inhibitory network in the cerebellar input layer, therefore has a number of important implications for cerebellar function.

## Acknowledgements

This project was supported by the Wellcome Trust (095667, 203048). R.A.S is in receipt of a Wellcome Trust Principal Research Fellowship. F.L. was supported by a postdoctoral Fondation Fyssen fellowship, a Marie Curie fellowship (FP7 program) and the Wellcome Trust (203048). We thank Lena Schmid and Antoine Valera for comments on the manuscript.

## Author contributions

F.L. and R.A.S. conceived the project and designed the experiments. F.L. and J.S.R performed the electrophysiological recordings, D.C. the immunohistochemistry and retrograde labelling. F.L. and R.A.S. wrote the manuscript.

## Declaration of Interests

None

## Methods

### Electrophysiology

#### Cerebellar slice preparation

Sagittal slices of the cerebellar vermis (230 μm thick), or coronal slices of the whole cerebellum (500 μm thick), were prepared from both male and female C57BL/6 mice (P22-33) in accordance with UK Home Office guidelines. Slices were prepared in a solution containing (in mM) 2.5 KCl, 4 MgCl_2_, 0.5 CaCl_2_, 1.25 NaH_2_PO_4_, 24 NaHCO_3_, 25 glucose, 230 sucrose, bubbled with 95% O_2_ and 5% CO_2_ and cooled to 2°C. Slices were immediately transferred into ACSF containing (in mM) 125 NaCl, 2.5 KCl, 2 CaCl_2_, 1 MgCl_2_, 1.25 NaH_2_PO_4_, 26 NaHCO_3_, and 25 glucose; pH = 7.3, equilibrated with 95% O_2_ and 5% CO_2_. Slices were maintained at 32°C for at least 30 min, then at room temperature for up to 4 h. Recordings were made at 32-36°C from cerebellar slices perfused in ACSF. Voltage-and current-clamp recordings from GoCs were performed with borosilicate glass pipettes (3 – 4 MΩ) containing (in mM) 140 KMeSO_4_, 6 NaCl, 1 MgCl_2_, 0.03 EGTA, 10 HEPES, 4 ATP-Mg, 0.4 GTP-Na, 10 Phosphocreatine and 6 Biocytin, unless otherwise stated. All chemicals were dissolved in water and obtained from Tocris Bioscience or Sigma-Aldrich.

#### Loose-cell-attached Golgi cell recordings

Spontaneous spiking activity in GoCs was recorded in a loose-cell-attached configuration with borosilicate patch pipettes (3 – 4 MΩ) containing ACSF. GoCs were selected based on their large somata in the GC layer and their spontaneous firing. Recordings were performed in the center of the GC layer, away from the Purkinje cell layer, in order to avoid recordings from Lugaro and globular cells (Hirono et al., 2012). Action potentials were identified based on prominent upward and downward deflections in the membrane current traces (Kanichay and Silver, 2008).

#### Golgi cell excitability

GoC R_input_ was calculated from steady-state membrane voltage responses to a 400 ms current pulse to −50 pA. The average spike frequency was calculated in response to 400 ms current pulses from 0 to 200 pA in steps of 10 pA. The slope was computed from a linear fit to the f/I relationship. The rheobase corresponds to the minimal amplitude of positive current injection necessary to trigger an action potential.

#### Golgi cell-Granule cell paired recordings

GoCs-GCs paired recordings were performed in ACSF containing 50 μM D-AP5 (D-(-)-2-Amino-5-phosphonopentanoic acid) and 10 μM NBQX (2,3-Dioxo-6-nitro-1,2,3,4-tetrahydrobenzo[*f*]quinoxaline-7-sulfonamide disodium salt) to isolate GC IPSCs. GoCs were held in current clamp and GCs were held in voltage clamp at −70 mV. GC IPSCs were recorded with the same high chloride internal solution used for recording spontaneous PSCs. 20 GoC action potentials were triggered at 20 Hz, repeated every 20 seconds, by injecting brief positive current pulses (1 nA, 1 ms). GC IPSCs were recorded during control, NE application and washout periods, averaged (n = 10-15 repetitions) and integrated to compute the charge transfer during the total IPSC and slow IPSC. The slow IPSC, reflecting the spillover, was estimated by clipping the fast component, and making the region between two successive stimuli equal to a linear fit taken between the average of the previous and last 20 ms of each interval. The fast IPSC component was determined by subtracting the slow IPSC from the total IPSC.

#### Glutamatergic synaptic transmission from parallel fibres and mossy fibres onto Golgi cells

Glass pipettes containing ACSF were placed in the molecular layer to stimulate PFs or in the white matter to stimulate MFs as described by Kanichay and Silver (Kanichay and Silver, 2008). 5 stimuli at 50 Hz were delivered every 20 seconds (Digitimer, 100 μs, 10 − 30 V). The stimulus intensity was set to 5 V above threshold. GoCs were voltage-clamped at −60 mV in the presence of 10 μM SR95531 (6-Imino-3-(4-methoxyphenyl)-1(6*H*)-pyridazinebutanoic acid hydrobromide) (gabazine) and 0.5 μM Strychnine to isolate alpha-amino-3-hydroxy-5-méthyl-4-isoxazolepropionic acid (AMPA) receptor-mediated currents. After a stable baseline recording of 5 min, NE (10 μM) was bath applied, followed by a washout. Stability of the whole-cell recordings was performed by monitoring pipette series resistance R_s_ as described above. Cells were discarded if R_s_ was > 25 MΩ or changed by > 20 % over time. GoC holding current was measured within a 100 ms window before the stimuli. EPSC peak amplitudes were baseline subtracted using a 0.5 ms window ~0.1 ms before the stimulus. The peak amplitudes were computed by averaging points within a 0.1 ms window centered on the peak. All mean peak amplitudes were calculated from the average of at least 30 repetitions. EPSC peak amplitudes were computed as described above.

#### Measuring the persistent sodium current in Golgi cells

Effects of NE on I_NaP_ in GoCs were measured using a slow depolarizing voltage ramp (−80 to −10 mV, slope 10 mV/s) before and after bath application of 10 μM NE. GoCs were patched via electrodes filled with an internal solution containing (in mM) 140 CsMeSO_4_, 4 NaCl, 2 MgCl_2_, 10 HEPES, 3 Na_2_ATP, 0.3 NaGTP, 5 Phosphocreatine and 0.2 EGTA, titrated to pH 7.3 with CsOH. 10 repetitions of the voltage-ramp protocol were averaged for each cell and for each condition in the presence of Gabazine (10 μM), Strychnine (0.5 μM), NBQX (10 μM) and D-AP5 (50 μM). Each average was baseline subtracted by fitting the linear portion between −80 and −70 mV to zero. The peak of the I_NaP_ was determined as the minimum point of a 7^th^ order polynomial fit applied to the baseline subtracted ramp between −65 mV and −10 mV. Recordings that showed excessive spiking activity during the ramp were excluded from the analysis.

### Data acquisition and analysis

Data was recorded using a MultiClamp 700B amplifier (Molecular Devices) and NeuroMatic software (Rothman and Silver, 2018) that works within the Igor Pro environment. Data analysis was performed using NeuroMatic or Prism (GraphPad). Data are presented as mean ± SEM of *n* experiments. Membrane potentials are specified without correction for the liquid junction potential. Voltage- and current-clamp recordings were low-pass filtered at 10 kHz and digitized at 20 kHz. All statistical analysis was performed in the Prism environment (GraphPad). Comparisons between groups were performed using non-parametric Wilcoxon matched pairs test. Significance levels were defined as p < 0.05.

### Surgical procedures

All surgical procedures were carried out in accordance with institutional animal welfare guidelines and licensed by the UK Home Office. Adult C57BL/6 mice (P40-60) were anesthetized with ketamine–xylazine, mounted in a stereotaxic frame and injected with 0.3 μl of undiluted red retrobeads IX (Lumofluor, USA) into lobule IV/V of the cerebellar vermis. After recovery, mice were then housed for 7 days before being sacrificed.

### Immunohistochemistry

Mice were deeply anaesthetised with sodium pentobarbital and transcardially perfused with 4% paraformaldehyde in phosphate buffered saline (PBS). Brains were extracted and post-fixed for 24 h. 60 µm coronal sections were prepared using a vibrating microtome (VT1000S, Leica Microsystems) and transferred to multiwell plates for free float immunolabelling. Nonspecific binding was blocked with 5% normal goat serum. Sections were incubated overnight with antibodies for tyrosine hydroxylase (1:700 dilution, Millipore AB152), then incubated with a fluorophore-conjugated secondary antibody (alexa 488 or alexa 568). Sections were slide mounted with vectashield antifade mounting medium and imaged with a Leica TCS SPE8 confocal microscope.

